# A single DNA binding site of DprA dimer is required to facilitate RecA filament nucleation

**DOI:** 10.1101/2024.05.07.592916

**Authors:** Irina Bakhlanova, Begoña Carrasco, Aleksandr Alekseev, Maria Yakunina, Natalia Morozova, Mikhail Khodorkovskii, Michael Petukhov, Dmitry Baitin

**Affiliations:** Petersburg Nuclear Physics Institute Named by B.P. Konstantinov of National Research Center «Kurchatov Institute», Gatchina, Russian Federation; Department of Microbial Biotechnology, Centro Nacional de Biotecnología, CNB-CSIC, 3 Darwin Str, 28049 Madrid, Spain; Peter the Great St. Petersburg Polytechnic University, St. Petersburg, Russian Federation

## Abstract

The DprA (*a.k.a*. Smf) protein has emerged as a RecA mediator during natural chromosomal transformation, but its ubiquity suggests a possible role in DNA metabolism beyond natural transformation. We show that *Bacillus subtilis dprA* increases the frequency of *Escherichia coli* Hfr conjugation. RecA·ATP binds and cooperatively polymerises in a 50-nucleotide (nt) poly deoxy T (dT)_50_ ssDNA to form a dynamic filament with SSB competing for binding, but *B. subtilis* DprA (DprA_*Bsu*_) counters the inhibitory effects of SSB on RecA·ATP filaments. RecA bound to (dT)_21_ is poorly active as dATPase, with DprA_*Bsu*_ significantly improving RecA dATP hydrolysis. RecA_*Bsu*_·dATP-(dT)_20_ complexes were readily formed, while DprA_*Bsu*_ exerts an allosteric effect on RecA_*Bsu*_-(dT)_15_ complexes competent for dATP hydrolysis. Combining experimental data with a full-atomic model of the RecA-DprA-ssDNA complex’s spatial structure, we proposed a molecular mechanism for DprA-mediated loading of RecA onto short ssDNA stretches. Our results suggest that steric constraints allow for the participation of only one DNA binding site of the DprA dimer in RecA-mediated dATP hydrolysis.

## 1. Introduction

DprA is an evolutionarily conserved and ubiquitous protein that directly interact with RecA and with the single-stranded binding proteins (SsbA and SsbB), as supported by yeast two-hybrid, pull-down and Förster resonance energy transfer data [1, 2]. The SsbA and/or SsbB proteins coated the incoming single-stranded (ss) DNA as soon as it exits from the entry channel. Dimeric DprA partially displaces the SsbA and/or SsbB proteins and loads RecA onto ssDNA to initiate natural chromosomal transformation [3]. RecA interacts with and destabilizes the DprA dimer interface resulting in ssDNA transfer from DprA to RecA [4]. DprA is also present in non-competent species, indicating its potential multifunctionality [5, 6]. The presence of DprA across bacterial species, coupled with a high degree of sequence conservation, suggests its vital functional role. DprA’s role as a natural chromosomal transformation factor in numerous Firmicutes species has been well-established, with extensive studies on DprA proteins from *Bacillus subtilis* (DrpA_*Bsu*_) and *Streptococcus pneumonia* (DrpA_*Spn*_) [2, 4, 7]. However, the primary function of DprA in non-naturally transforming bacteria remains elusive. DprA binds to cytosolic SsbA/SsbB-ssDNA, playing a crucial role in RecA-dependent natural chromosomal transformation and RecA-independent natural plasmid transformation [7, 8]. Unless stated otherwise, the genes and products mentioned are of *E. coli* origin. Proteins derived from other organisms are designated using an abbreviation that includes the *Genus* and species of the respective organism (*e.g*., *B. subtilis* DprA is referred to as DprA_*Bsu*_).

The ability of DprA from various bacterial species to bind both ssDNA and dsDNA has been demonstrated [2, 7, 9]. The DprA has at least three discrete biochemical activities: (a) it partially displaces SsbA or SsbB negative mediators and interacts with RecA and loads onto ssDNA, as a positive mediator [7]; (b) it catalyses strand annealing to the complementary ssDNA strand coated by the SsbA or SsbB protein [8]; and (c) it counteracts the negative effect of the negative RecX modulator on RecA filament growth [10]. Additionally, DprA_*Spn*_ triggers competence shut-off to avoid a detrimental consequence of prolonged natural competence [11]. *In vitro* reconstituted assays revealed that DprA_*Spn*_ displaces SSB from ssDNA and facilitates RecA nucleation and filament growth on the DprA_*Spn*_-ssDNA-SSB complexes [2], suggesting an heterospecific interaction between evolutionarily distant DprA_*Spn*_ and SSB and RecA proteins.

DprA has been identified in many bacterial species not known to undergo natural competence development, leading to the hypothesis that DprA functions beyond transformation in these bacteria. The *dprA* (*a.k.a. smf*) gene product has the ability to restore natural chromosomal transformation in *Haemophilus influenza* competent cells [12]. This suggests an *in vivo* heterospecific interaction between evolutionarily conserved DprA and RecA_*Hin*_ proteins, highlighting its biological significance. The DprA is a 374-amino acid residue protein. While the crystal structure of full-length DprA is not available, the structure of DprA_*Spn*_ has been elucidated [13]. The DprA_*Spn*_ structure consists of the association of a sterile alpha motif domain and a Rossmann fold, forming tail-to-tail dimers. Key interaction residues positioning on *the* DprA_*Spn*_ structure indicates an overlap of DprA_*Spn*_-DprA_*Spn*_ and DprA_*Spn*_-RecA_*Spn*_ interaction surfaces [13]. Overall, the structural features of DprA_*Spn*_ protein suggest a competitive interaction between DprA_*Spn*_ and RecA_*Spn*_ lacking the first 27 residues [4]. The multiple pieces of evidence of heterospecific interaction between DprA_*Spn*_ and RecA have been previously obtained *in vitro*, suggesting a possible non-cognate protein for use as a tool [2]. To determine whether a Firmicutes DprA is capable of functioning during Hfr conjugation we investigated the role of *B. subtilis* DprA (DprA_*Bsu*_) protein when expressed from a plasmid. DprA_*Bsu*_ was selected because full-size DprA_*Bsu*_ and RecA_*Bsu*_ was detected yeast two-hybrid assays, whereas a binary interaction between full-size DprA_*Spn*_ and RecA_*Spn*_ was not detected in a yeast two-hybrid assay [13]. Indeed, a RecA_*Spn*_ variant lacking the N-terminal monomer-monomer interacting domain, interacts with full-size DprA_*Spn*_ [13].

The DprA_*Bsu*_ protein when expressed from a plasmid. By controlling the time of protein induction and temperature incubation, we demonstrate an almost twenty-fold increase in FRE. Additionally, we tested the effect of *dprA*_*Bsu*_ on Hfr conjugation for comparative purposes, as it is well-characterized and shares a significant identity and similarity (42% and 62%, respectively) with DprA.

There are no published accounts that the DprA protein has been purified for biochemical studies. The DprA is highly susceptible to proteolytic breakdown and is generally insoluble in aqueous solutions. In the current study, we demonstrate that DprA_*Bsu*_ partially displaces SSB from ssDNA and facilitates RecA nucleation and filament growth by indirect measurement RecA-mediated ATP hydrolysis. The detection of heterospecific interactions between DprA_*Bsu*_ and RecA, together with the similar degree of conservation observed between RecA_*Bsu*_ and RecA, led us to use RecA to investigate the effect of DprA_*Bsu*_ binding to small oligodeoxythymidylate, (dT), of defined length. RecA, in the ATP bound form (RecA·ATP), nucleated on a (dT)_34_ oligo is not active as ATPase, but the presence of DprA_*Bsu*_ rendered RecA·ATP competent in ATP hydrolysis. dATP induces a very high affinity ssDNA binding state of RecA [14]. RecA·dATP is poorly active as ATPase on (dT)_21_, but in the presence of DprA_*Bsu*_ was competent in ATP hydrolysis. To further evaluate such protein-protein interaction we have purified and used homologous RecA_*Bsu*_. We also propose a full-atomic model of the spatial structure of the RecA-ssDNA-DprA complex build using molecular modelling. The model describes sterically possible form of the RecA–ssDNA-DprA complex capable to explain the experimental data.

## 2. Result

### 2.1. DprA significantly increases the frequency of recombinational exchanges (FRE)

Constant presentation of new ssDNA occurs during conjugation, as the donor DNA is transferred into recipients [15]. Throughout mating, the donor Hfr KL227 transfers markers into recipients sequentially in the order of *leu*^+^, *ara*^+^, and *thr*^+^. The recombination exchanges in this region of the *E. coli* map are adequately described mathematically by the Haldane formula. Over the past 20 years, a method for measuring recombination exchanges to assess elevated or depressed levels of recombination in vivo has been developed. This method has the advantage of detecting essentially all types of genetic exchanges that might occur during conjugation [16].

At least two quantitative parameters have been utilized to characterize conjugational recombination - the yield of recombinants and the linkage of two transferred donor markers. The frequency of recombinational exchanges is expressed as FRE, the average number of exchanges per one *E. coli* genome equivalent (100 min). For wild-type *E. coli*, FRE = 5.0. The FRE approach aligns with methodologies developed by others [17, 18]. The ΔFRE is thus FRE1/FRE2, where FRE1 is simply the FRE measured for the cross under investigation. We explored a potential role for *dprA* in conjugation through Hfr conjugation experiments, utilizing *dprA* or *dprA*_*Bsu*_. The genes encoding the DprA protein or DprA_*Bsu*_ protein were expressed in the presence of IPTG. We have adjusted the time of protein overexpression to avoid the artefact of metabolic overload due to protein overproduction and to get closer to more physiological levels for genetic studies. The differential effects of the two DprA proteins are readily observed in FRE values under normal conditions. There was a 2.6 and 6.4-fold increase in FRE values, relative to the empty vector, in the transconjugants observed in cells expressing DprA or DprA_*Bsu*_. The data presented in this section indicate that, compared to DprA, the DprA_*Bsu*_ protein appears to be a stronger regulator of HR activity, measured through FRE, the stimulatory role of which was measured at 37 °C (Table 1a). To facilitate the accumulation of a database that is more optimally matched for protein activity, we conducted the following studies at a 30 °C temperature of protein induction. Surprisingly, the FRE level increased for both proteins. In effect, the 30 °C condition of protein induction led to a 17-to 19-fold increase in FRE values (Table 1b). The DprA proteins contributed equally to the recombination potential under conditions of protein over-expression at 30 °C.

**Table 1a.**
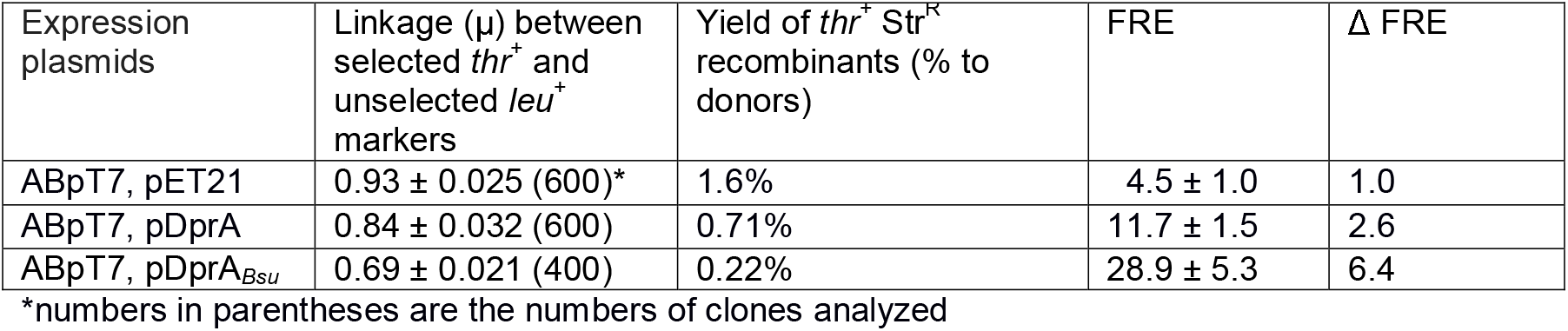
The dependence of the FRE value on the increased expression of DprA in transconjugants of the AB1157 line crossed with the KL227 donor, at 37 °C.

**Table 1b.** The dependence of the FRE value on the increased expression of DprA in transconjugants of the AB1157 line crossed with the KL227 donor, at 30 °C.

| Expression plasmids | Linkage ($\mu$) between selected *thr*<sup>+</sup> and unselected *leu*<sup>+</sup> markers | Yield of *thr*<sup>+</sup> Str<sup>R</sup> recombinants (% to donors) | FRE | $\Delta$ FRE |
| --- | --- | --- | --- | --- |
| ABpT7, pET21 | 0.949±0.024 (600)* | 1.7% | 3.2 ± 0.6 | 1.0 |
| ABΔdprA, pET21 | 0.94 ± 0.018 (600) | 3.1% | 3.6 ± 0.5 | 1.1 |
| ABpT7, pDprA | 0.59 ± 0.025 (800) | 0.2% | 55.5 ± 6.3 | 17.3 |
| ABpT7, pDprA<sub>BSU</sub> | 0.56 ± 0.036 (600) | 0.02% | 61.2 ± 8.5 | 19.1 |

\*numbers in parentheses are the numbers of clones analyzed

As previously described, normal FRE values for the wild-type background appear to be induced in a temperature-dependent manner, showing a 2-fold increase in FRE at 42 °C, respectively [16]. This suggests that temperatures as high as 37 °C inhibit the intracellular amount or activity of proteins or both. This result leads to two major conclusions. The primary conclusion is that the DprA protein can be viewed as an important regulator of HR in Gram-negative bacterial conjugation with a strong stimulatory effect. Second, DprA_*Bsu*_ efficiently replaces DprA in *E. coli* cells, showing a similar FRE value, regardless of the mechanism of action behind the positive effect.

### 2.2. DprA promotes SSB dislodging from ssDNA

Previously is has been shown that: i) in the presence of mediators RecA efficiently nucleates and polymerizes on the ssDNA-SSB complexes, and RecA-mediated ATP hydrolysis is used as an indirect inference of RecA dynamic assembly on the ssDNA-SSB complexes; and ii) DprA is a genuine mediator. To test whether DprA_*Bsu*_ promotes the nucleation of RecA protein filaments onto SSB-coated ssDNA, the circular ssDNA (12 μM) was pre-incubated with SSB protein was sufficient to bind all DNA present in the reaction -0.5 μM of protein. Then, increasing DprA_*Bsu*_ and sub-stoichiometric amount of RecA (1 μM) were added and the steady state rate of RecA-mediated ATP hydrolysis measured. The reaction exhibited active displacement of SSB protein, evidenced by an increase in ATPase with a rise in the concentration of the regulatory protein DprA (Figure 1a).

**Fig. 1.**
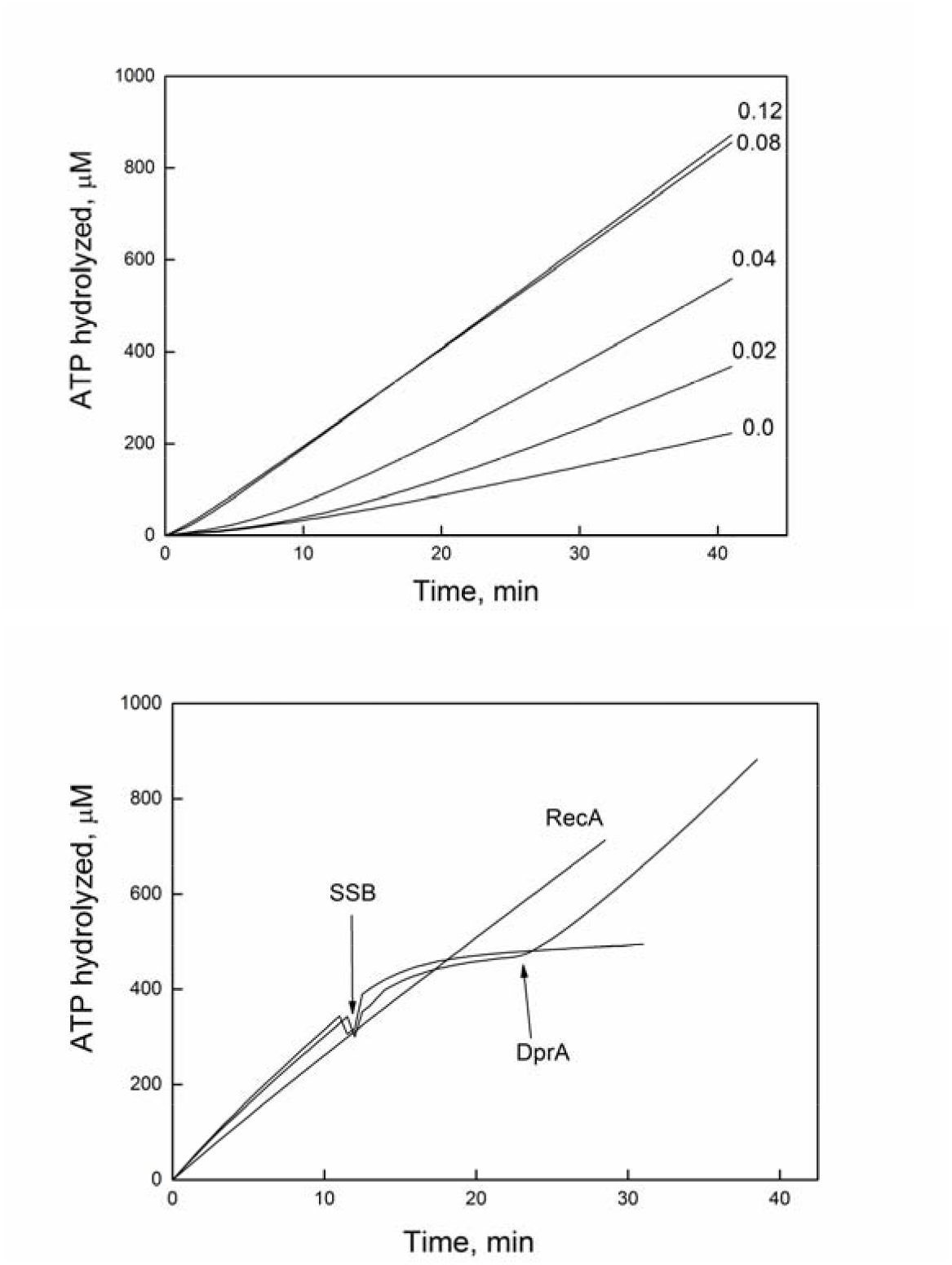
*DprA_Bsu_ antagonises SSB function. (a) The reactions mixture contained 12 μM M13mp8 ssDNA and 0.5 μM SSB. Then 1 μM RecA and various DprA_Bsu_ protein concentration as indicated. b)* DprA_*Bsu*_ can reverse the effects of SSB protein on RecA-mediated ATP hydrolysis. *The reactions were carried out as described under “Experimental Procedures” and contained 3 μM RecA protein, 5 μM (dT)_50_ (length of the oligo) All experiments were carried out at least three times with consistent results*.

The result suggests that DprA_*Bsu*_ effectively stimulates RecA under analogous conditions and concentration ratios as observed in the referenced study. In the initial minutes, a lag phase of ATP hydrolysis was observed, indicating a temporary delay in filament nucleation due to the obstacle presented by SSB. Subsequently, the ATP hydrolysis kinetics reached a steady-state, indirectly reflecting the proportion of each participating protein remaining bound to DNA. The maximum rate of RecA-mediated ATP hydrolysis is achieved in the presence of 1 DprA_*Bsu*_ monomer per 150-nt of DNA. Hence, the impact of heterospecific protein interactions, considering the provided concentrations, aligns with data on homospecific interactions published earlier. Ultimately, these findings affirm the existence of heterospecific interactions between *RecA* and DprA_*Bsu*_ proteins in vitro, signifying the evolutionary conservation of protein-protein interactions.

The quantitative equilibrium between SSB and RecA proteins at a steady state during their competitive interplay is affected not only by their concentrations and order in which they are added to the reaction mixture, but also by the nuances of the DNA architecture. If, on a circular DNA of any length, RecA tends to displace SSB, then on linear molecules with a free 5’ end, conversely, SSB displaces RecA, as the disassembly of the RecA filament progresses in the 5’→3’ direction [19, 20]. As the filaments dissociate from the ends, the time of disassembly depends on the number of ends. The shorter the linear DNA molecule, the shorter filament would dissociate in a shorter time and hence promptly replaced by the SSB protein. The kinetic outcomes of RecA filament disassembly are predictably different when long ssDNAs are substituted by ssDNA stretches. Natural DNA molecules, such as long phage DNA, exhibit diverse secondary structures hindering filament polymerization. Known for unravelling DNA hairpins, the SSB protein aids the filament in extending along the DNA.

Each of these regulatory proteins may indirectly influence filamentation efficiency through DNA structure, adding complexity to data interpretation. In this particular experiment, a (dT_n_) oligo was used to eliminate potential side effects associated with circular single-stranded DNA, focusing on the direct effects of protein-protein interactions involving DprA_*Bsu*_, SSB, and RecA. The RecA concentration this experiment is ample, even in excess, to bind the entire (dT_50_) oligonucleotide. As illustrated, SSB promptly, almost instantaneously, displaced RecA from the (dT_50_), consistent with previously known data (Figure 1b). However, addition of DprA_*Bsu*_ completely facilitated formation of an active RecA filament, as the ATP hydrolysis rate in its presence remains unchanged following the addition of SSB. The observed steady state ATP hydrolysis in the presence of DprA_*Bsu*_ remains constant, unaffected by the order of protein addition. Even if the (dT_50_) is already bound by SSB protein, the kinetics are reversible. The newly established steady-state ATP hydrolysis is indistinguishable from the initial one. Given the absence of secondary structure on (dT_50_), all effects can be attributed to direct protein-protein interactions. Within this experiment, the sequence of SSB protein replacement by other proteins remains ambiguous. One mechanism was that DprA initiates binding first, displacing SSB protein by interaction and providing an available platform on DNA for RecA assembly [7, 21]. Another possible mechanism entails RecA and DprA_*Bsu*_ both engage with DNA cooperatively to increase the event of nucleation as if RecA alone were acting in a concentration dependent manner.

These two mechanisms are not mutually exclusive. The second mechanism, which ascribes greater significance to the interaction between RecA and DprA_*Bsu*_, is implied by the considerably enhanced capability of DprA_*Bsu*_ to initiate RecA filamentation on the DNA.

### 2.3. DprA_*Bsu*_ facilitates RecA nucleation onto short *(dT*_*n*_*) oligos*

To examine the effect of DprA_*Bsu*_ on the ability of RecA nucleation and filament growth, RecA was pre-incubated with oligos of variable length, followed by the addition of DprA_*Bsu*_. As previously documented, the affinity of the RecA filament for the ssDNA substrate increased with the length of the ssDNA oligo, resulting in the formation of a highly stable ssDNA-RecA complex with (dT)_50_ oligo [22, 23]. Varying the length of the substrate helps to isolate the nucleation phase from filament extension. Given the cooperative nature of monomer binding, an insufficient number of subunits in the filament compromises the complex’s stability. The other current view is that the protein nucleates as an oligomer of 5 subunits or even as a dimer, forming a compact cluster on the DNA [24, 25]. Anyway, in fact, a short (dT)_30_ to (dT)_40_ oligo proves to be a poor substrate for filament formation. The interaction of the protein with DNA of such limited length is accompanied by only modest ATP hydrolysis. However, as the DNA length increases, the RecA-mediated hydrolysis rate gradually rises, reaching its peak values. A (dT)_34_ oligo can accommodate 10-11 RecA monomers. Considering some dynamic equilibrium during the dissociation and association of the complex, the actual number may be even lower at any point of time. At the concentration of RecA protein used in our experiment, ATP hydrolysis is activated by less than 20% of the potential maximum value (Figure 2a). Thus, the potential for RecA nucleation and filament growth is severely limited. The addition of 0.06 μM DprA_*Bsu*_ dramatically elevates the rate of RecA-mediated ATP hydrolysis, regardless of the order of protein addition to the DNA. On the linear segment of RecA-dependent ATPase enhancement, it occurs proportionally to the amount of DprA_*Bsu*_ protein added. In fact, saturation occurs in the presence of one DprA_*Bsu*_ dimer/ (dT)_34_ (0.24 μM DprA_*Bsu*_). Taken together with the structural data, these results imply that the DprA_*Bsu*_ protein must interact with each DNA oligo for full nucleation stimulation. In other words, it must interact with every single filament associated with (dT)_34_.

**Fig. 2.**
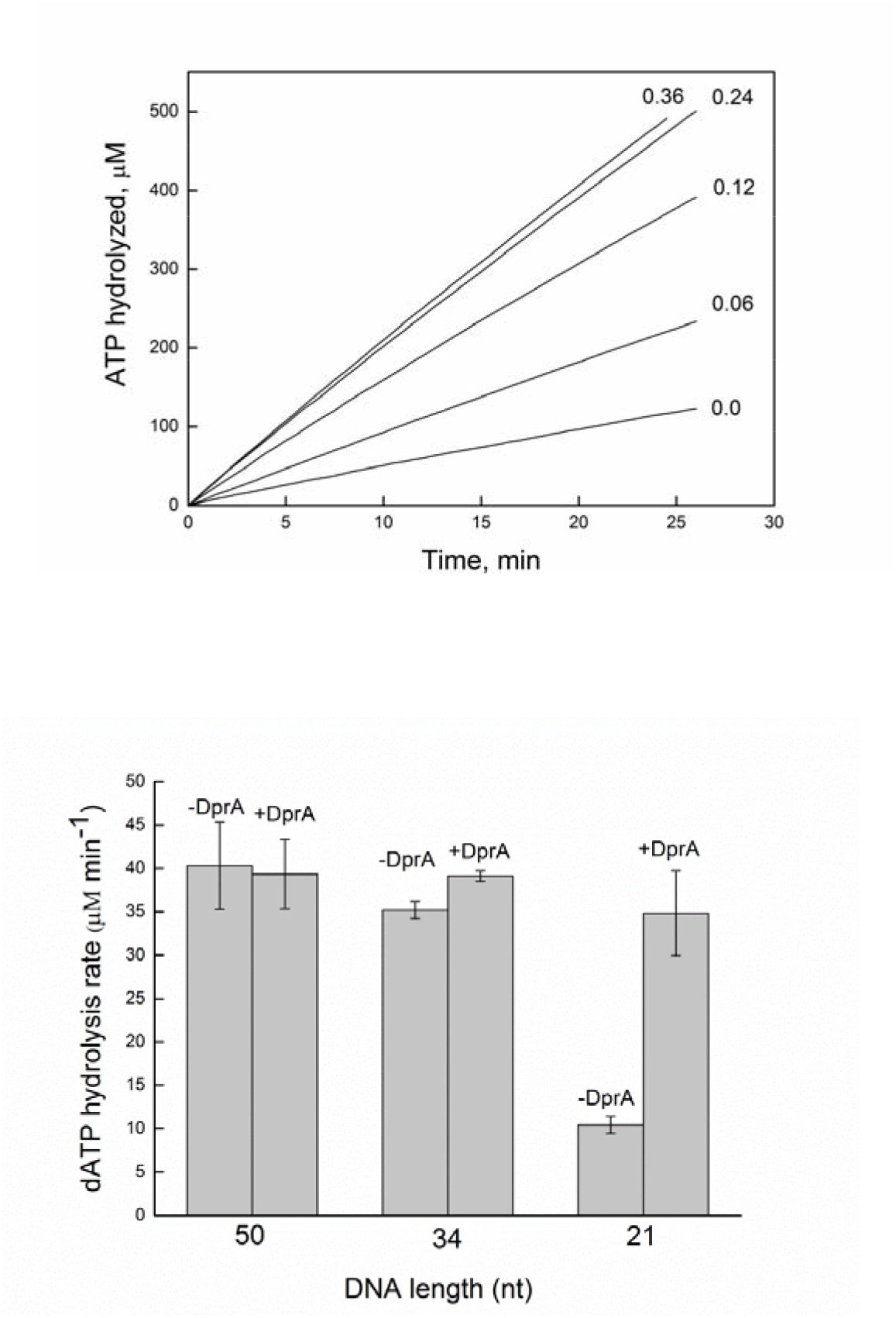
RecA protein can nucleate on the short ssDNA stretches in the presence of DprA_*Bsu*_. (a) *The reactions were carried out as described under “Experimental Procedures” and contained 3 μM RecA protein, 5 μM (dT)_34_ (in nucleotides) and 2 mM ATP. All experiments were carried out at least three times with consistent results. (b)* Effect of DNA size on RecA-dependent ATP hydrolysis in the presence or absence of DprA_*Bsu*_ protein. *The reactions were carried out as described under “Experimental Procedures” and contained 5 μM RecA protein, 0.3 μM DprA_Bsu_, 5 μM (dT)_n_, 2 mM dATP. The error bars are one standard deviation from the mean calculated from three independent experiments*

DprA_*Bsu*_ may interact with and facilitate the loading of RecA onto two types of substrates: those of suitable length for filament formation and those of shorter length. To more precisely determine the minimum substrate length allowing any stimulation of RecA nucleation by DprA_*Bsu*_, we refined the experimental conditions. Previous studies indicated that in the presence of dATP, filament disassembly slows down, and nucleation events on DNA are more frequent [14, 26]. Adjustments were made to enhance RecA binding affinity for short oligos, removing filament length as a limiting factor in the RecA-DprA_*Bsu*_ pair. This was achieved by substituting ATP for dATP in the hydrolysis assay, that is more consistent with the preferences of RecA.

Switching from ATP to dATP was therefore rationalised to determine the minimum site size required to accommodate the DprA_*Bsu*_-RecA complex. RecA·dATP bound to (dT)_34_ is competent in ATP hydrolysis (Figure 2b). To assess the minimum ssDNA length required for stable interactions with RecA, dATPase assays were performed using (dT)_50_ to (dT)_21_ ssDNA, with and without DprA_*Bsu*_ (Figure 2b). The (dT)_50_ ssDNA surpasses the critical threshold for sufficient RecA nucleation, rendering the contribution of DprA_*Bsu*_ to the transition of RecA from a free state to a bound state visually as if undetectable. Although DprA_*Bsu*_ remains present in the formation of long RecA filament, as indicated by the SSB protein displacement experiment, its stimulatory role becomes more evident with shorter oligonucleotides, increasing with oligo shortening and, consequently, filament length reduction.

The RecA·dATP bound (dT)_21_ was significantly impaired in dATP hydrolysis, but competent to hydrolysed dATP when bound to (dT)_34_ regardless of the presence of DprA_*Bsu*_. These data have demonstrated DprA’s ability to specifically promote the loading of RecA·dATP onto substrates as short as (dT)_21_. Obviously, the (dT)_21_ ssDNA, which corresponds approximately to the size of seven RecA protomer, is competent to activate the dATPase of RecA in the presence of 0.3 μM DprA_*Bsu*_ (Figure 2b).

### 2.4. *DprA*_*Bsu*_ promotes the formation of dynamic RecA_*Bsu*_·dATP filaments on (dT)_20_ ssDNA

To determine the minimum substrate length required for DprA to stimulate RecA nucleation, we optimized the experimental conditions using homologous proteins and dATP. As previously reported, RecA_*Bsu*_ is inactive as a dATPase in the absence of ssDNA, and DprA_*Bsu*_ cannot hydrolysed dATP [8]. This indicates that the dATP hydrolysis observed in the assays can be solely attributed to the nucleation of RecA_*Bsu*_·dATP onto short stretches of ssDNA.

To explore the mechanism by which DprA_*Bsu*_ promotes RecA_*Bsu*_ filament loading, we measured dATP hydrolysis. RecA_*Bsu*_·dATP (1 RecA monomer/ 2-nt (dT)_n_) binds to and cooperatively polymerises on (dT)_30_ ssDNA, forming a dynamic filament that hydrolysed dATP with a k_cat_ of 2.3 ± 0.3 min^-1^ (Figure 3). In the presence of (dT)_20_ and (dT)_15_ ssDNA, RecA_*Bsu*_·dATP exhibited dATP hydrolysis rates with k_cat_ values of 0.2 ± 0.02 min^-1^ and 0.19 ± 0.02 min^-1^, respectively, slightly above the background levels observed with RecA or DprA in the absence of ssDNA.

**Figure 3.**
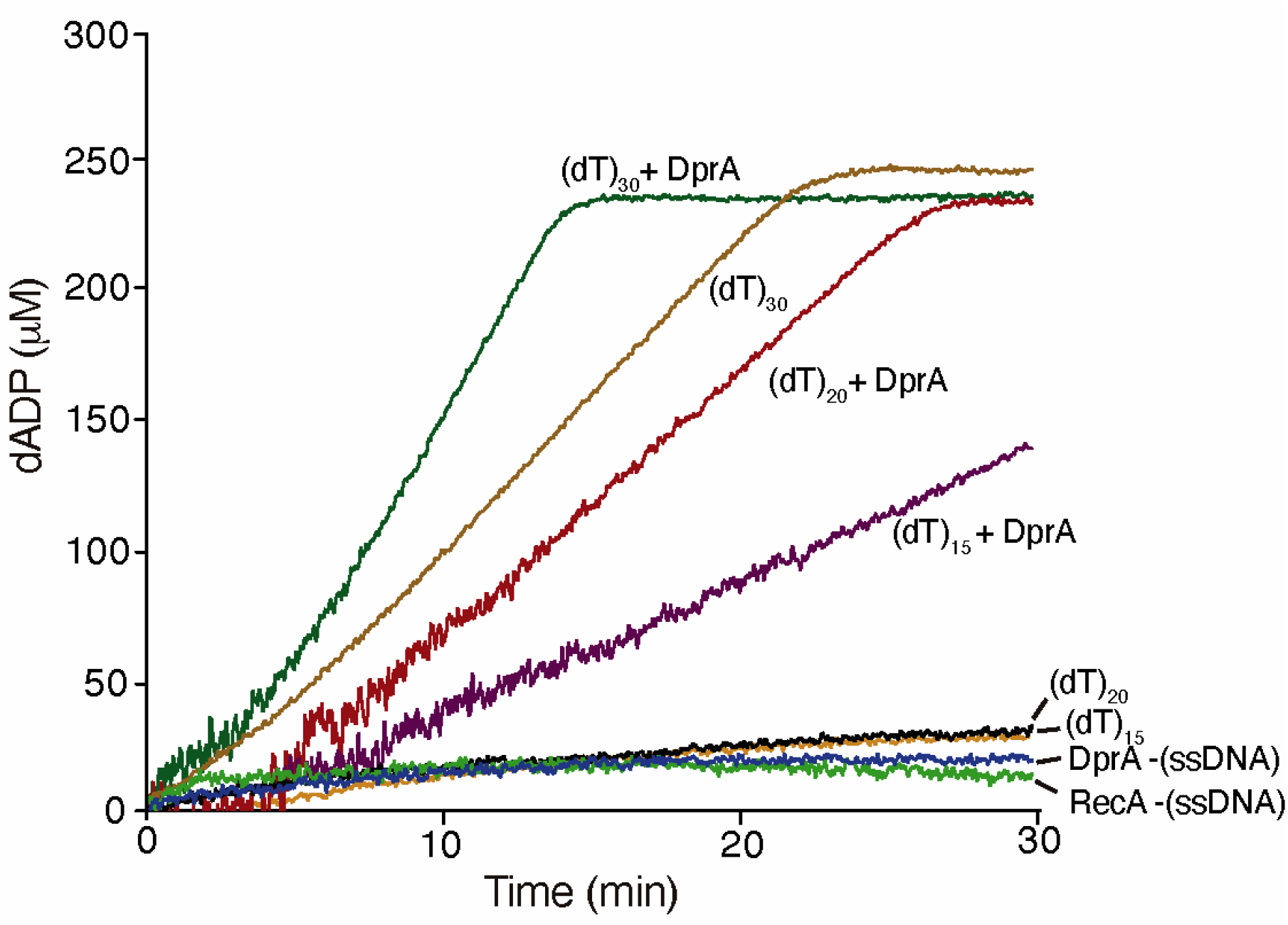
Effect of DNA size on RecA_*Bsu*_-dependent dATP hydrolysis in the presence or absence of DprA_*Bsu*_ protein. *Reactions were performed as described in the “Experimental Procedures” section and contained 3 μM RecA_Bsu_ protein, 10 μM (dT)_n_, 2.5 mM dATP in Buffer B*. When indicated DprA_*Bsu*_ (*0.3* μM) was added to the reaction mixture where indicated. *Standard deviations were calculated from the mean of three independent experiments*.

To test whether RecA_*Bsu*_·dATP undergoes a functional transition upon interaction with DprA_*Bsu*_, and whether short DNA substrates might confer structural stability for RecA loading, DprA_*Bsu*_ was added to the reaction mixture. Under these conditions, RecA_*Bsu*_ nucleated and polymerised on (dT)_30_ ssDNA, forming a dynamic filament that rapidly hydrolysed dATP with a k_cat_ of 4.4 ± 0.4 min^-1^. The presence of DprA_*Bsu*_ not only stabilised but also enhanced the final rate of RecA_*Bsu*_-mediated dATP hydrolysis when bound to (dT)_20_ or (dT)_15_, yielding k_cat_ values of 2.0 ± 0.4 min^-1^ and 1.1 ± 0.2 min^-1^, respectively (Figure 3).

The co-crystal structures showed 5-nt binding per DrpA_*Hpy*_ monomers and 3-nt per RecA monomer [27, 28], and DprA_*Bsu*_ and RecA_*Bsu*_ physically interact (see above). Therefore, we reasoned that DprA_*Bsu*_ bound to (dT)_20_ ssDNA interacts with and loads RecA_*Bsu*_·dATP, contributing to the loading of ∼5 RecA_*Bsu*_·dATP protomers. The DprA_*Bsu*_-RecA_*Bsu*_ interaction may elicit an allosteric effect on both proteins. Under this condition monomeric DprA binds 5-nt ssDNA and RecA_*Bsu*_ cooperativity increased facilitating dynamic filaments. This is consistent with the observation that a DprA_*Spn*_ monomer has only a 2.6-fold weaker binding affinity for (dT)_20_ compared to the DprA_*Spn*_ dimer [13]. DprA_*Spn*_ or DprA_*Bsu*_ is sufficient to recruit homologous or heterologous RecA, in the ATP or dATP bound form, onto short ssDNA oligos [2, 7, 8], hence for simplicity we will discuss the interaction of DprA on the RecA-ssDNA complexes.

### 2.5. Reconstruction of the structure of the RecA-DprA-ssDNA complex

Our RecA·dATPase and RecA_*Bsu*_·dATPase experiments reveals a further reduction of the binding site of DprA_*Bsu*_. DprA_*Bsu*_ interacts with and caps and stabilises RecA/RecA_*Bsu*_ filament on a (dT)_21/20_ ssDNA. However, with only biochemical experiments involving ATPase or dATPase analyses and no structural studies, the picture is not clear. A DprA_*Bsu*_-RecA_*Bsu*_·dATP complex bound to a (dT)_20_ or (dT)_15_ ssDNA substrate enhances dATP hydrolysis throughout the filament and yields RecA_*Bsu*_·dADP. Then, RecA_*Bsu*_·dADP protomer-protomer interaction within the filament stabilises DNA binding, with dissociation occurring predominantly from filament ends. In other words, the interaction of DprA_*Bsu*_ and RecA_*Bsu*_ should be sufficient to allosterically stabilise RecA_*Bsu*_ to cooperatively bind ssDNA to form helical nucleoprotein filaments, with its N-terminal helix destabilising a DprA_*Bsu*_ dimer in favour of a functional monomer, as early suggested [4, 13]. This would provide a molecular explanation for the transfer of short (dT) ssDNA from DprA_*Bsu*_ to RecA_*Bsu*_. This hypothesis inspired us to refine an existing model in which the structure of the complex could fit the experimental data.

The figure 4 shows a molecular model of the DprA_2_-RecA_3_·ATP-(dT)_19_ complex constructed on the basis of available structural data for three different complexes, which data on the spatial structure are published [13, 27, 28]. Although this nucleoprotein complex has a multicomponent structure, obtained from the structural data of complexes of various compositions, the structure and conformation of its constituent proteins turned out to be compatible with each other and does not have any structural problems. However, the conformation of the 5’-end of the ssDNA presynaptic filament RecA (shown in brown) turned out to be incompatible with the presence of DprA proteins and directed towards the region of DprA protein-protein interface, and not towards the known nucleotide binding site found in the crystal structure of the dimeric complex of DprA proteins with short fragments of ssDNA [28]. This obviously means that binding of DprA proteins requires that the six nucleotide residues of the free 5’-end of ssDNA adopt a suitable conformation for binding to the nucleotide binding site of the DprA protein. Molecular modelling of the conformational transition of ssDNA showed that the nucleotide from the dimeric complex of the DprA protein [28] and ssDNA of the presynaptic RecA protein filament [27] can be connected without any steric strains by two mobile nucleotide residues, thus forming a continuous ssDNA chain running from the nucleotide binding site of the DprA protein to the RecA proteins helical filament as shown in the figure 4.

**Figure 4.**
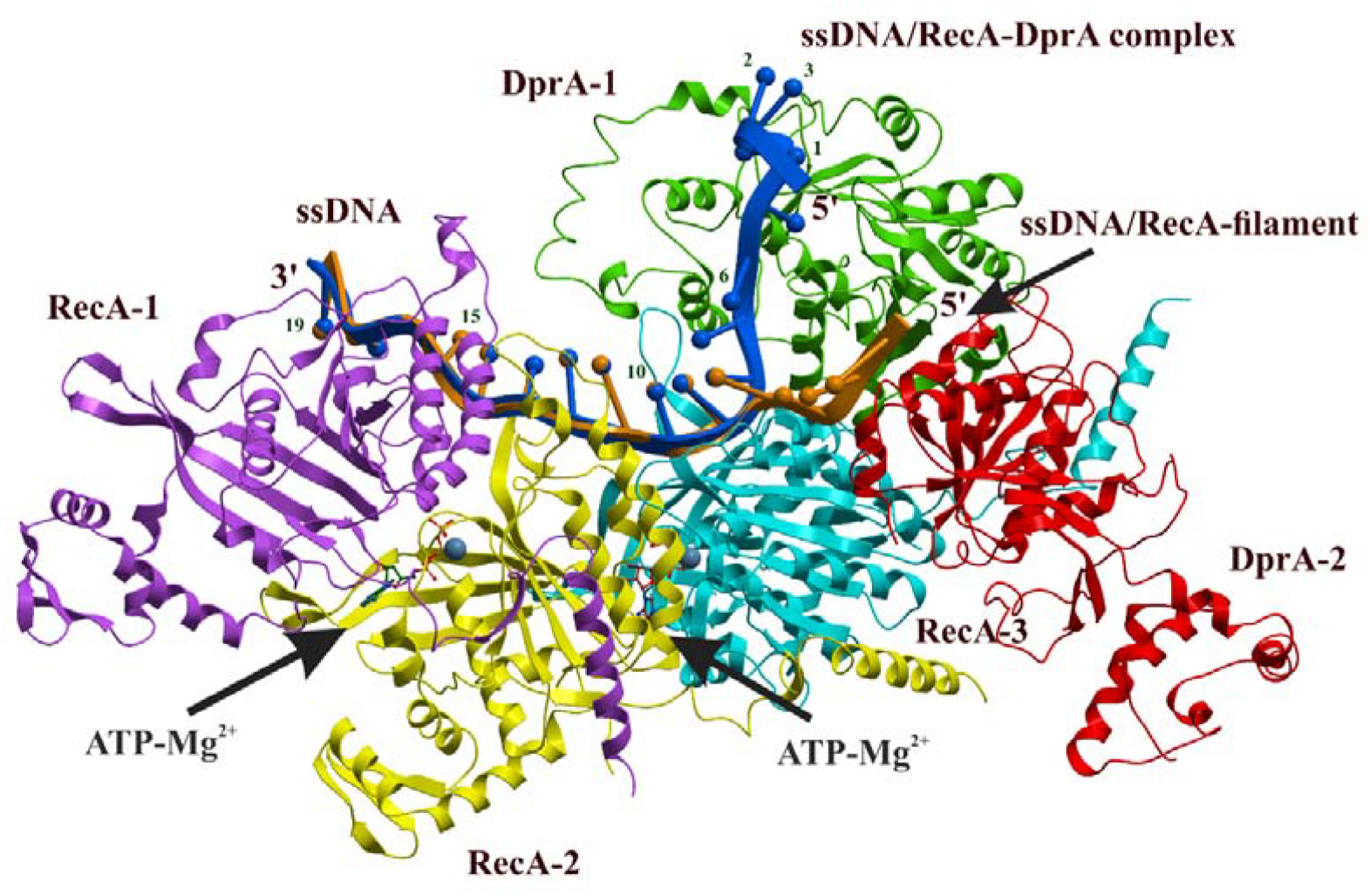
The molecular model of the complex of the RecA nucleoprotein filament (DprA_2_-RecA_3_·ATP-(dT)_19_. Dimeric DprA, with protomers shown in green and red, and the three RecA protomers are shown in blue, yellow and magenta respectively. The conformation of the ssDNA (dT_19_) where the conformation of the first six 5’ terminal nucleotide residues is taken from ssDNA-DprA complex (PDB ID: 4LJR) [13] while the conformation of the last eleven 3’ terminal nucleotide residues is taken from the crystal structure of the presynaptic RecA nucleoprotein filament (PDB ID: 3CMX) [27] is shown in blue. The original conformation of the ssDNA from the presynaptic RecA nucleoprotein filament is shown in brown for clarity. Two ATP–Mg^2+^ substrate complexes in the RecA protein ATPase sites are shown in ball and stick representation. (PDB see in Supplemental Data)

The ATP/dATP hydrolysis site is located at the interface between two protomers of RecA. While the binding of RecA monomers to ssDNA in the filament structure requires the formation of direct contacts between loops L1 and L2 of neighbouring protomers every three nucleotide residues of ssDNA. Thus the attachment of each new subunit of the RecA protein to the nucleoprotein filament requires three nucleotide residues of ssDNA [27].Therefore, the minimum length of ssDNA required for all standard intermolecular interactions of a DprA-RecA complex to be formed for at least one ATP binding site is 15-nt as depicted in Figure 4. In the case of two ATP binding sites (three RecA proteins), which certainly provides a higher rate of ATP hydrolysis, the minimum sufficient length of ssDNA in a DprA-RecA complex is 18-20 nt. As shown in Figures 2 and 3, the experimental results are consistent with the theoretical model of the complex.

One of the interesting questions is the role of the second subunit of the DprA protein in the complex with DprA_2_-RecA_3_·ATP-(dT)_19_ shown in Figure 4. It is obvious that exactly the same symmetrical structure can be built on the basis of the complex of the second subunit of DprA with RecA protein. However, the simultaneous existence of these two structures in one complex is sterically impossible due to the competition of two symmetrically located subunits of the RecA protein for the same place in space.

Based on recently obtained structural data and the results of molecular modelling of complexes of the DprA protein with DNA and the RecA protein, a molecular mechanism of DprA-mediated loading of RecA proteins onto ssDNA has been proposed [29]. This molecular mechanism involves the formation of a dimeric nucleoprotein complex of DprA with ssDNA, which provides a suitable platform for further nucleation of the RecA filament, after which one of the DprA subunits leaves the complex and merization of RecA on ssDNA continues. It is assumed in the model that ssDNA binds simultaneously to two DprA subunits, the DNA binding sites of which are located at a distance of approximately 50Å, which, according to measurement done on the molecular model, corresponds to the length of ssDNA having 10 nucleotide residues in a fully unfolded extended conformation. Experimental analysis of the binding of ssDNA of various lengths to oligomeric complexes of DprA proteins showed that the stability of their binding depends on the DNA length [28]. Short fragments of ssDNA having 6–20 nucleotides in length either do not bind to DprA at all or bind with very low affinity. High-affinity complexes were obtained for ssDNA having 35 nucleotides in length. The structure of the dimeric complex of DprA proteins with DNA was obtained using X-ray methods. However, most of the (dT)_35_ ssDNA turned out to be disordered, and the electron density of ssDNA was detected only for the 6-nt of the 5’ end. The conformation of these 6-nt were used to reconstruct the DprA_2_-RecA_3_·ATP-(dT)_19_ complex shown in Figure 4. RecA interacts with and destabilizes the DprA dimer interface resulting in ssDNA transfer from DprA to RecA.

It is also important that due to the symmetry of the dimeric complex of the DprA protein, the ssDNA chains in the ssDNA binding sites of the DprA proteins were directed in opposite directions. This makes it impossible to form a dimeric complex of DprA with one strand of ssDNA occupying both sites of the DprA proteins. Therefore, it is sterically possible to form of only a complex of DprA with ssDNA, occupying only one DprA protein ssDNA binding site as shown in Figure 3. As for the role of the second subunit of the DprA, its role is apparently limited to participation in the mechanism of loading the first terminal monomer of RecA protein onto ssDNA. The second subunit also has an effect on the affinity of the first subunit DprA, for ssDNA. And in this case, the minimum length of ssDNA is much shorter and amounts to 6+11=17-nt residues. This estimate of the minimum required length of ssDNA explains the experimental data obtained in this work.

## 3. Discussion

We investigated the functionality of the DprA homolog in *E. coli* in the context of Hfr conjugation. While most regulators only partially alter the genetic outcomes of recombination, thereby adjusting the natural FRE values used to assess *in vivo* recombinase activity [16, 18, 30, 31], our study reveals a striking observation under normal conditions: DprA/Smf emerges as a potent activator of recombination activity *in vivo*, significantly augmenting FRE upon overexpression (by approximately 18-fold). It is conceivable, that the DprA–RecA effect in *E. coli* is likely to be regarded as an additional modulation of the main recombination pathways, the exploration of which will contribute to a more comprehensive understanding of homologous recombination. Taken together, these studies demonstrate that DprA proteins can play different roles in different bacteria.

The second statement is based on the clarifying experiments, the main one being the search for the minimum length of the substrate capable of providing a platform for the nucleation of the RecA filament. The fact is that the real length of DNA required for a complex containing both proteins is only about twenty nucleotides. This result inspired us to refine an existing model in which the structure of the complex could fit our experimental data. Our study is based on the preexisting analysis based on mutagenesis data of key amino acid residues that enabling DprA protein to bind DNA and RecA [4]. This study confirmed that the dimeric form of DprA is required for effective DNA binding and loading RecA recombinase. Similarly, the biochemical characterisation of a monomeric, dimerisation-deficient DprA mutant from *H. pylori* showed that only the dimeric form is functionally active in DNA binding [28]. Our experimental data and theoretical modelling provide a new molecular mechanism to explain the previously shown model with the same key determinants. The structural and biochemical characterisation of the DprA protein reported here showed that it is sterically possible to form only a complex of DprA_2_ with ssDNA where DprA occupies only one binding site with ssDNA. In the complex, the second DprA subunit is thought to sterically stabilise the first RecA monomer in DNA binding form. Given the known size of the DprA-DNA binding site and dimer DrpA as active species, we extrapolate these data to construct a ternary RecA -DprA -ssDNA complex that occurs during the RecA filament nucleation. Within this context, the remaining 21 nt ssDNA olig space can accommodate, at most, a RecA trimer as the nucleation unit, which subsequently fills the rest of the substrate upon DprA dissociation, forming a hexamer. The complex structure between DprA and ssDNA reported here provides us with a good starting point for further studies on DprA and other related proteins. Undoubtedly, regulators of homologous recombination involved in the processes of chromosomal transformation should be considered as potential targets for inhibition [32, 33]. Further study of the structure of the DprA-RecA complex may lead to the design of compounds or inhibitors of horizontal gene transfer.

## 4. Material and Methods

### 4.1 Strains and plasmids

Donor KL227 (HfrP4x metB) and recipients: AB1157 (*thr-1 leuB6 ara14 proA2 hisG4 argE3 thi-1 supE44 rpsL31*) and recombination deficient JC10289 (as AB1157 but Δ*recA-srlR306::Tn10* [*=* Δ*recA306*]) were from A.J. Clark’s collection. Strain JW5708-1 (Δ*dprA-724::kan*) was from Keio Collection (*E. coli* Genetic Resources at Yale CGSC). A null *dprA* (Δ*dprA*) strain was constructed by P1 transduction to transfer *dprA*-724 (del)::kan from strain JW5708-1 into AB1157 as described previously [34]. Plasmid pT7 (original name pT7POL26) codes for T7 RNA polymerase under the control of a *lacZ* promoter. This plasmid was used to express DprA/Smf proteins under conditions of lac promoter induction by IPTG. Plasmids pDprA_*Bsu*_ was constructed and kindly supplied by Prof. J.C. Alonso. Plasmid pDprA was constructed and supplied by Eurogen. These plasmids contain the DprA and DprA_*Bsu*_ genes under a T7 RNA polymerase driven promoter.

### 4.2 Proteins

The wild-type RecA and RecA_*Bsu*_ were purified as previously described [35, 36]. Single-strand binding (SSB) protein was kindly provided by Prof. M. Cox (University of Wisconsin-Madison) Their concentrations were determined using native extinction coefficients: _j_280_ = 2.23 × 10^4^ M^−1^cm^−1^ for RecA [37], 1.52 10^4^ M^−1^cm^−1^ for RecA_*Bsu*_ and 2.38 × 10^4^ M^−1^cm^−1^ for SSB protein [38]. The concentration of DprA _*Bsu*_ was determined using the native extinction coefficient _j_280_ = 4.5 × 10^4^ M^−1^cm^−1^ [7].

DprA_*Bsu*_ was purified similarly to the previously described protocol with minor modifications [7]. Briefly, BL21(DE3) pLysE cells were transformed with the pCB888 plasmid kindly provided by prof. J.C. Alonso. Bacterial cells were grown at 25°C and expression was induced by the addition of 0.4 mM isopropyl-1-thio-β-D-galactopyranoside (IPTG) at an OD600 of 0.5. After induction, cells were grown for 3 hours and rifampicin (200 µg/ml) was added 30 minutes after IPTG addition. DprA was purified from the clarified lysate by two-step chromatography using HisTrap HP 1 mL (GE Healthcare) and HiTrap SP XL 1 mL columns (GE Healthcare). Fractions containing DprA were concentrated on an Amicon Ultra-4 centrifugal filter (Merck) with a 3 kDa cut-off, supplemented with 50% glycerol and stored at -20°C.

### 4.3. (d)ATP hydrolysis Assays

A coupled enzyme, spectrophotometric assay [39, 40] was used to measure RecA-mediated ATP or dATP hydrolysis. The ADP or dADP generated by hydrolysis was converted back to ATP or dATP by a regeneration system of pyruvate kinase and phosphoenolpyruvate (PEP). The resultant pyruvate was converted to lactate by the lactate dehydrogenase using NADH as a reducing agent. The conversion of NADH to NAD^+^ was monitored as a decrease in absorbance at 380 nm. The amount of ATP or dATP hydrolysed over time was calculated using the NADH extinction coefficient ε_380_ = 1.21 mM^−1^cm^−1^. The assays were carried out in a Cary 5000 dual beam spectrometer at 37°C, with a temperature controller and a 12-position cell changer. The path length was 1 cm, the band pass was 2 nm. All RecA ATPase or dATPase assays were performed in buffer A (25 mM Tris-HCl (pH 7.5, 88% cation), 10 mM MgCl_2_, 5% w/v glycerol, 1 mM dithiothreitol (DTT), 2 mM ATP or dATP) containing 3 mM PEP, 10 U/ml pyruvate kinase, 10 U/ml lactate dehydrogenase, 4.5 mM NADH and 5 μM M13mp18 cssDNA or (dT) oligos. Whereas, RecA_*Bsu*_ dATPase assays were performed in buffer B (50 mM Tris-HCl [pH 7.5], 10 mM Mg(OAc)_2_, 5% w/v glycerol, 1 mM dithiothreitol (DTT), 2.5 mM dATP) containing 0.5 mM PEP, 10 U/ml pyruvate kinase, 10 U/ml lactate dehydrogenase, 4.5 mM NADH and (dT)_n_ oligos (10 μM). All reactions were repeated at least three times with similar results.

### 4.4. Conjugation

Conjugation was carried out essentially as described [16]. Both Hfr and F−strains were grown, crossed and selected for recombinants at 37°C or 30°C as indicated in the table, for 1 hour, in mineral salts 56/2 medium supplied with all necessary growth factors at pH 7.5. The ratio between donors and recipients in the mating mixture was 1:10, 2–4 x107 donors and 2–4 × 108 recipients per 1 ml. The yield of *thr*^+^ Str^r^ and *ara*^+^ Str^r^ recombinants in all independent crosses (5–7% relative to donors) was normalized according to the mating ability of each recipient used. The latter was determined by the yield of transconjugants F’-lac^+^ in crosses between the recipients and donor P200 F’-lac.

FRE value calculations were carried out as described [16]. Alterations in FRE (ΔFRE) promoted by the mutant *recA* gene or by any accessory gene relative to the FRE value promoted by the wild type Ec-*recA* gene were calculated using the following formula: ΔFRE = ln(2μ1 - 1)/(2 μ2 - 1), where μ1 is the linkage of selected thr+ or ara+ and unselected leu+ markers in a cross using wild type E. coli strain AB1157 and μ2 is the similar linkage in the cross being analysed. Calculations of uncertainty of relative FRE values were determined as deviations from the average values by making use of the program Excel-97 with formula [=2*STDEV] and by inputting the values from independent repeats of three experiments.

### 4.5. Reconstruction of the structure of the RecA-ssDNA-DprA_*Bsu*_ complex

To construct a full-atomic model of the spatial structure of the RecA-ssDNA-DprA_*Bsu*_ complex, we used a set of crystal structures from the Protein Data Bank, namely the structure of the presynaptic nucleoprotein filament of the RecA protein containing RecA_6_–(ADP–AlF_4_–Mg)_6_–dT_18_ (PDB ID: 3CMX) [27], the structure of *Helicobacter pylori* DprA_*hpy*_ protein with ssDNA (DprA_*Hpy*_–dT_6_)_2_ (PDB ID: 4LJR) [28] and the spatial model of the RecA-DprA complex obtained using molecular docking methods, which was kindly provided by the authors of this work [13]. The structure superimposition, energy minimization and other molecular manipulations were carried out using standard protocols of the ICM-Pro software package (Molsoft LLC, USA), [41]) and the ECEPP/3 force field as implemented in the ICM-Pro [41, 42].

The spatial structure of the dimeric complex of the DprA proteins from *E. coli* was obtained based on the crystal structure of the DprA_*hpy*_ dimeric complex (PDB ID: 4LJR) using standard homology modelling protocols implemented in ICM-Pro. The RecA-sDNA-DprA_*Bsu*_ complex was assembled using a series of superimpositions of the spatial structure of proteins. At the first stage, the position of the DprA_*Bsu*_ dimer was determined using the superimposition of RecA monomers in the presynaptic nucleoprotein filament (PDB ID: 3CMX) [27] and in the spatial model of the RecA-DprA_*Bsu*_ complex [13]. The position of 6-nt at the 5’-end of ssDNA was then determined using superposition of the DprA_*Hpy*_-dT_6_)_2_ complex (PDB ID: 4LJR) [28] and the molecular model of the RecA-DprA complex, while the conformation of the 3’-terminal part of ssDNA was taken in the form in which it is located in the presynaptic nucleoprotein filament RecA (PDB ID: 3CMX) [27]. Finally, the conformation of 2 bridging nucleotides was determined by minimizing the energy of ssDNA with fixed 3’ and 5’ ssDNA ends.

## Acknowledgements

We are grateful to Prof. Juan C. Alonso (Department of Microbial Biotechnology, Centro Nacional de Biotecnología, Madrid, Spain) for his generous gift of expression plasmids bearing the B. subtilis DprA for the E. coli strain. The authors would like to thank Prof. Michael M. Cox (University of Wisconsin– Madison) for SSB protein. The authors thank Jessica Andreani and Sophie Quevillon-Cheruel (Institute of Integrative Biology of the Cell, Université Paris-Sud, Gif-sur-Yvette, France) for structural model of the complex of the DprA dimer with RecA protein. We acknowledge the help of Inna Kurdyumova, Daria Antonova and Maryam Dmitrieva (Peter the Great St. Petersburg technic University) in purification of *B. subtilis* DprA. B.C. acknowledges financial support from Ministerio de Ciencia e Innovación MCIN/Agencia Estatal de Investigación (AEI)/ 10.13039/501100011033/ FEDER, EU, PID2021-122273NB-I00, as well as from CSIC 2021AEP031 and 202520E100.

